# Identification of Antigenic Regions Responsible for inducing Type 1 diabetes mellitus

**DOI:** 10.1101/2022.07.20.500753

**Authors:** Nishant Kumar, Sumeet Patiyal, Shubham Choudhury, Ritu Tomer, Anjali Dhall, Gajendra P. S. Raghava

## Abstract

There are a number of antigens that induce autoimmune response against β-cells, leading to Type 1 diabetes mellitus (T1DM). Recently several antigen-specific immunotherapies have been developed to treat T1DM. Thus identification of T1DM associated peptides with antigenic regions or epitopes is important for peptide based-therapeutics (e.g., immunotherapeutic). In this study, for the first time an attempt has been made to develop a method for predicting, designing and scanning of T1DM associated peptides with high precision. We analyzed 815 T1DM associated peptides and observed that these peptides are not associated with a specific class of HLA alleles. Thus, HLA binder prediction methods are not suitable for predicting T1DM associated peptides. Firstly, we developed a similarity/alignment based method using BLAST and achieved a high probability of correct hits with poor coverage. Secondly, we developed an alignment free method using machine learning techniques and got maximum AUROC 0.89 using dipeptide composition. Finally, we developed a hybrid method that combines the strength of both alignment free and alignment based methods and achieve maximum AUROC 0.95 with MCC 0.81 on independent dataset. We developed a webserver “DMPPred” and standalone server, for predicting, designing and scanning of T1DM associated peptides (https://webs.iiitd.edu.in/raghava/dmppred/).

**Key Points:**

- Prediction of peptides responsible for inducing immune system against β-cells
- Compilation and analysis of Type 1 diabetes associated HLA binders
- BLAST based similarity search against Type 1diabetes associated peptides
- Alignment free method using machine learning techniques and composition
- A hybrid method using alignment free and alignment based approach

**Author’s Biography:**

1. Nishant Kumar is currently working as Ph.D. in Computational biology from Department of Computational Biology, Indraprastha Institute of Information Technology, New Delhi, India
2. Sumeet Patiyal is currently working as Ph.D. in Computational biology from Department of Computational Biology, Indraprastha Institute of Information Technology, New Delhi, India
3. Shubham Choudhury is currently working as Ph.D. in Computational biology from Department of Computational Biology, Indraprastha Institute of Information Technology, New Delhi, India
4. Ritu Tomer is currently working as Ph.D. in Computational biology from Department of Computational Biology, Indraprastha Institute of Information Technology, New Delhi, India
5. Anjali Dhall is currently working as Ph.D. in Computational Biology from Department of Computational Biology, Indraprastha Institute of Information Technology, New Delhi, India.
6. Gajendra P. S. Raghava is currently working as Professor and Head of Department of Computational Biology, Indraprastha Institute of Information Technology, New Delhi, India.

## Introduction

Diabetes mellitus (DM) is a chronic, metabolic disorder majorly occurring due to the abnormality in the blood sugar or glucose level, which further leads to serious damages in heart, eyes, nerves, kidney and blood vessels [1, 2]. According to the WHO (world health organization) around 422 million people in the world are effected with DM and approximately 1.5 million deaths have been reported every year. DM is majorly categorized into two types i.e., T1DM (Type 1 diabetes mellitus) or insulin dependent diabetes and T2DM (Type 2 diabetes mellitus) or adult onset diabetes [1, 3]. T2DM affects middle-aged and older persons who suffer from persistent hyperglycaemia and affects around 6.28% of the world’s population and also known as a lifestyle disorder [4]. On the other side, T1DM is an immune-mediated serious lifelong incurable autoimmune disorder that affects around 5-10% of all cases of diabetes and mainly identified in children or adolescents. It is a serious condition in which insulin producing β-cells in the pancreas are destroyed [5, 6]; insulin is a peptide hormone that regulates the blood glucose levels [7, 8]. Yoon *et. al*. reports that factors which involve in the pathogenesis of autoimmune diabetes includes macrophages, T-lymphocytes, B-lymphocytes, β-cell autoantigens and dendritic cells. [9].

The pathogenesis of T1DM includes both genetic as well as environmental factors [10] [See Figure 1]. It has been observed in some studies that environmental factors play a significant role in disease development by triggering islet autoimmunity [11]. The genetic factors involved in the disease severity are mainly human leukocyte antigen (HLA) class-I/II and expression in β-cells [12]. Previous studies also report that the presence of specific HLA-alleles (HLA-DRB1*04/DQB1*0302/DRB1*03), autoantibodies, autoreactive and anti-islet antigen-specific T-cells are mostly identified in the T1DM patients [13-17]. As shown in Figure 1, several diseases are associated with the progression of T1DM such as celiac disease, coronary heart disease, ratinopathy, rheumatoid arthritis, autoimmune thyroid diseases, type A gastritis, vitiligo, systemic lupus erythematosus, and Addison disease, etc. [18-21].

**Figure 1:**
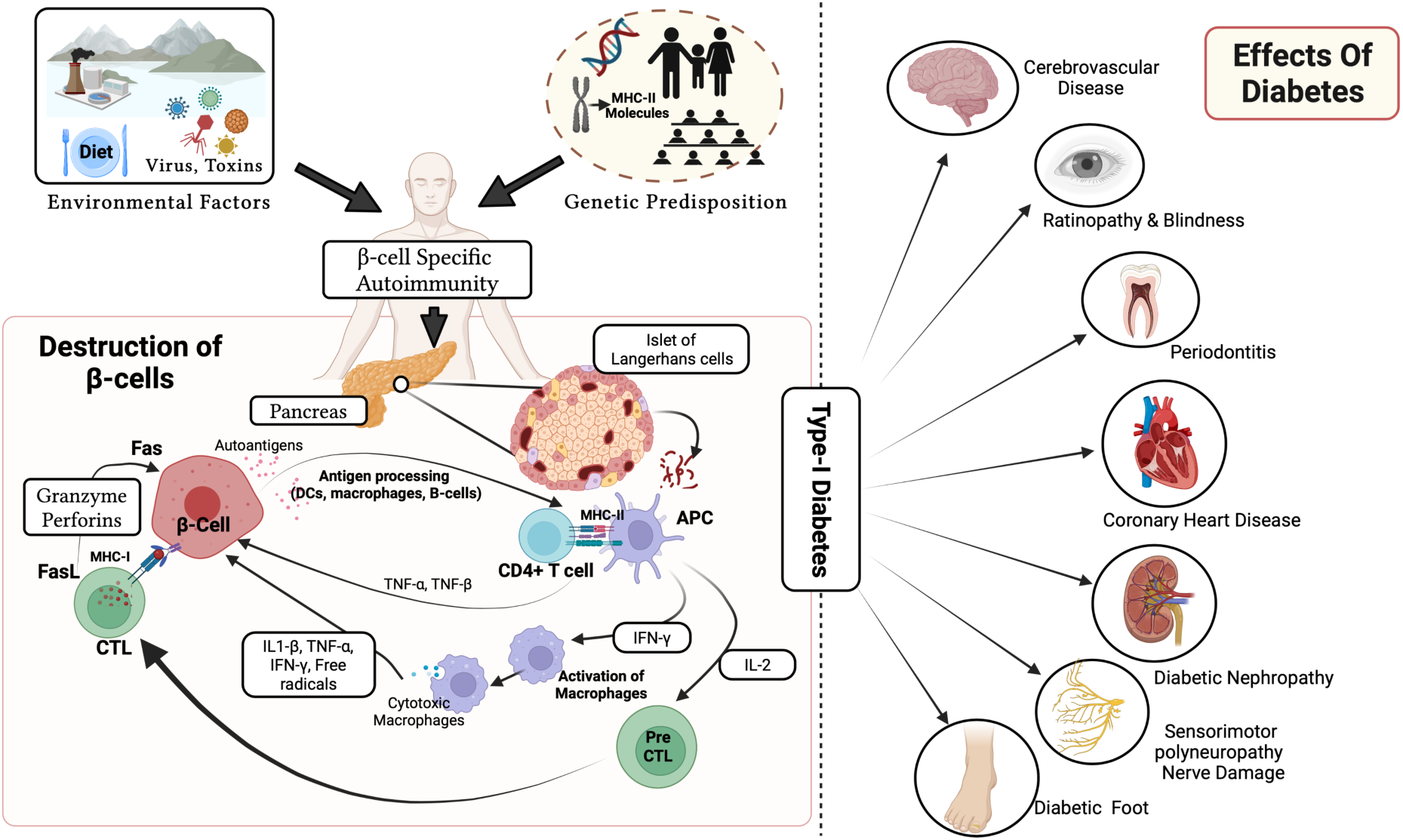
Detailed mechanism of β-cells destruction and various effects of diabetes.

Identification of T1DM associated peptides is an crucial step in understanding the fate of disease. These peptides mainly bind to MHC class II alleles and induce immune responses responsible for destroying β-cells. In the previous studies, limited attempts have been made to predict T1DM associated peptides using *in silico* techniques. Cia et al, developed a method GPS-MBA 1.0 for the prediction of T1DM associated MHC class II binders. The major limitation of this study is that it is developed on a small dataset consisting of only HLA-DQ8 binders. Thus, it is the need of the hour to develop a better and accurate tool for the prediction of T1DM associated peptides using latest available information. In this study, we have collected experimentally validated T1DM associated peptides from the immune epitope database (IEDB) [22]. Due to the limited availability of negative data, we generated random peptides from Swiss-Prot [23]. We analysed these peptides to understand their characteristics.

## Material and methods

### Dataset Collection

In order to construct a successful prediction model, data acquisition is the primary and cumbersome step [24, 25]. The IEDB is a major resource in the field of immunology, as it maintains experimentally validated MHC binders and epitopes [25]. We extracted T1DM associated peptides from IEDB and, all redundant peptides were removed. It was observed that most of the peptides have 8 to 30 amino acids, so we removed peptides having length more than 30 amino acids or less than 8 amino acids. Finally, we got 815 unique T1DM associated peptides which we labelled as positive data. Due to the limited availability of experimentally validated non-T1DM associated peptides, we generate random peptides from proteins in Swiss-Prot [26, 27]. Finally, we got 815 T1DM associated peptides and 815 random (non-T1DM associated) peptides. In order to evaluate models without any biasness, we used 80% of data for building models called training datasets. Remaining 20% data is used for testing our models, this is called independent/validation dataset. Our independent dataset is not used for any training or tuning hyperparameters, it is only used for evaluating the final model. On the other hand, our training dataset is used for training and testing for optimizing variables of models.

### Composition Analysis

Amino acid composition (AAC) is the compositional representation of peptide sequences, which represents the percent occurrence frequency of 20 amino acids in the protein/peptide sequence. It generates a 20-dimensional feature vector that specifies the number of each type of amino acid normalized with the total number of amino acids in the length of a provided peptide sequence, and can be calculated by using the following equation: [24, 25]

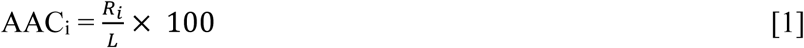

Where AAC_i_ is the percent composition of an amino acid i ; R_i_ is the number of residues of type i, and L is the total number of residues in the peptide [27].

### Sequence Logo

For the generation of sequence logos we have used the software named ‘Weblogo’. It provides the graphical representation of stack of amino acids. The sequence conservation is measured in bits. The logo generation requires FASTA or CLUSTAL formats. The overall height of each stack indicates the sequence conservation at that position, whereas the height of symbols within the stack reflects the relative frequency of the corresponding amino or nucleic acid at that position [28].

### Feature Generation

In order to build an alignment free model, we need to generate features or descriptors corresponding to peptides. As the length of peptides varies from 8 to 30 amino acids, we need to compute composition based features [24, 29]. In this study, we compute a wide range of compositional features like Amino acid composition (AAC), Dipeptide composition (DPC), Atom composition (ATC) using Pfeature [30].

### Feature Selection

Another crucial step in classification is feature selection, as most of the features are not significant. It is commonly used to identify the most relevant features. Pfeature generates a large pool of features and [31] mostly redundant and irrelevant features exist in the original feature set, which can create over-fitting [29]. The selection of the best features can minimize the risk of over-fitting and increase efficiency by reducing the model’s complexity [29, 32]. Selecting the relevant set of features from the enormous dimension of features is one of the primary concerns in this study. There are numerous approaches for feature selection [26]; we adopted mRMR (Minimum redundancy maximum relevance) algorithm[32].

### Machine learning techniques

We used a number of machine learning approaches to construct prediction models [27] including Decision Tree (DT), Logistic Regression (LR), Gaussian Naive Bayes (GNB), Random Forest (RF), k-nearest neighbor (KNNs), Extra-Trees (ET), XGBoost (XGB) and Support Vector Classification (SVC). The scikit-learn python library was used to implement these classification approaches [26].

### Five-Fold Cross-Validation

In order to develop general models that are not biased/overfitted for a dataset, we have implemented five-fold cross-validation algorithm [26, 31]. According to standard protocols of five-fold cross-validation the complete training dataset was divided into five equal sets. Where the four sets were designated as training dataset and fifth set assigned as testing dataset. In the results section, we presented the average scores of five cycles after repeating this approach five times.

### Similarity search

One of the commonly used techniques for annotation of proteins/peptides is based on similarity search. In this technique, the query peptide is aligned with all peptides whose function is known. The query peptide is annotated based on its alignment score with known peptides. One of the commonly used method for similarity search is BLAST [33-36]. Currently, we have implemented BLAST based search for the identification of similarity of peptides/epitopes with T1DM causing and non-T1DM causing peptides. This module is created using blastp (BLAST+ 2.7.1), a peptide-peptide BLAST which returns the most similar sequence from the database for the query sequence. Initially, a database have been created using the sequences in the training dataset, and query sequences from the independent dataset were hit against it at various e-values ranging from 1e-6 to 1e+4. We have considered the top-hits only and assigned the class based on the same, such as, if the top-hit is positive then the query sequence was assigned as positive and vice-versa.

### Evaluation parameters

Our study includes the well-established evaluation parameters for the evaluation of the machine learning models. We include both the parameters i.e. threshold dependent parameters and threshold-independent parameters. Sensitivity, Specificity, Accuracy, and Matthew’s correlation coefficient (MCC) are threshold dependent parameters while Area Under the Receiver Operating Characteristic (AUROC) curve are threshold-independent parameter [26, 27, 31]. These measurements are calculated using the following equations (2-6).

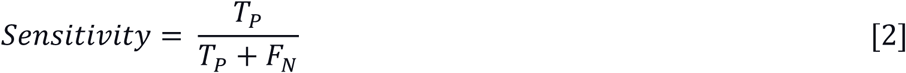

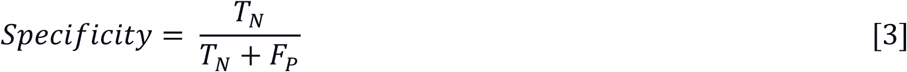

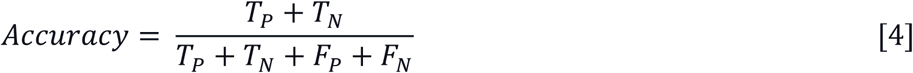

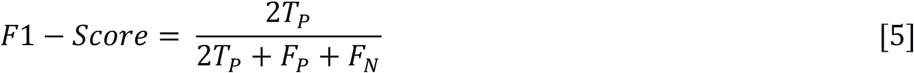

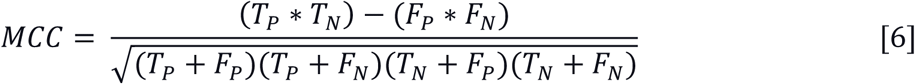

Where, T_P_, T_N_, F_P_ and F_N_ stands for true positive, true negative, false positive and false negative, respectively.

## Results

In this study, we incorporate 815 T1DM associated peptides and we call it as positive dataset. The negative dataset includes random 815 non-T1DM associated peptides generated from proteins in Swiss-Prot. The prediction and analysis was performed on these peptides, Figure 2 represents the overall workflow of the study.

**Figure 2:**
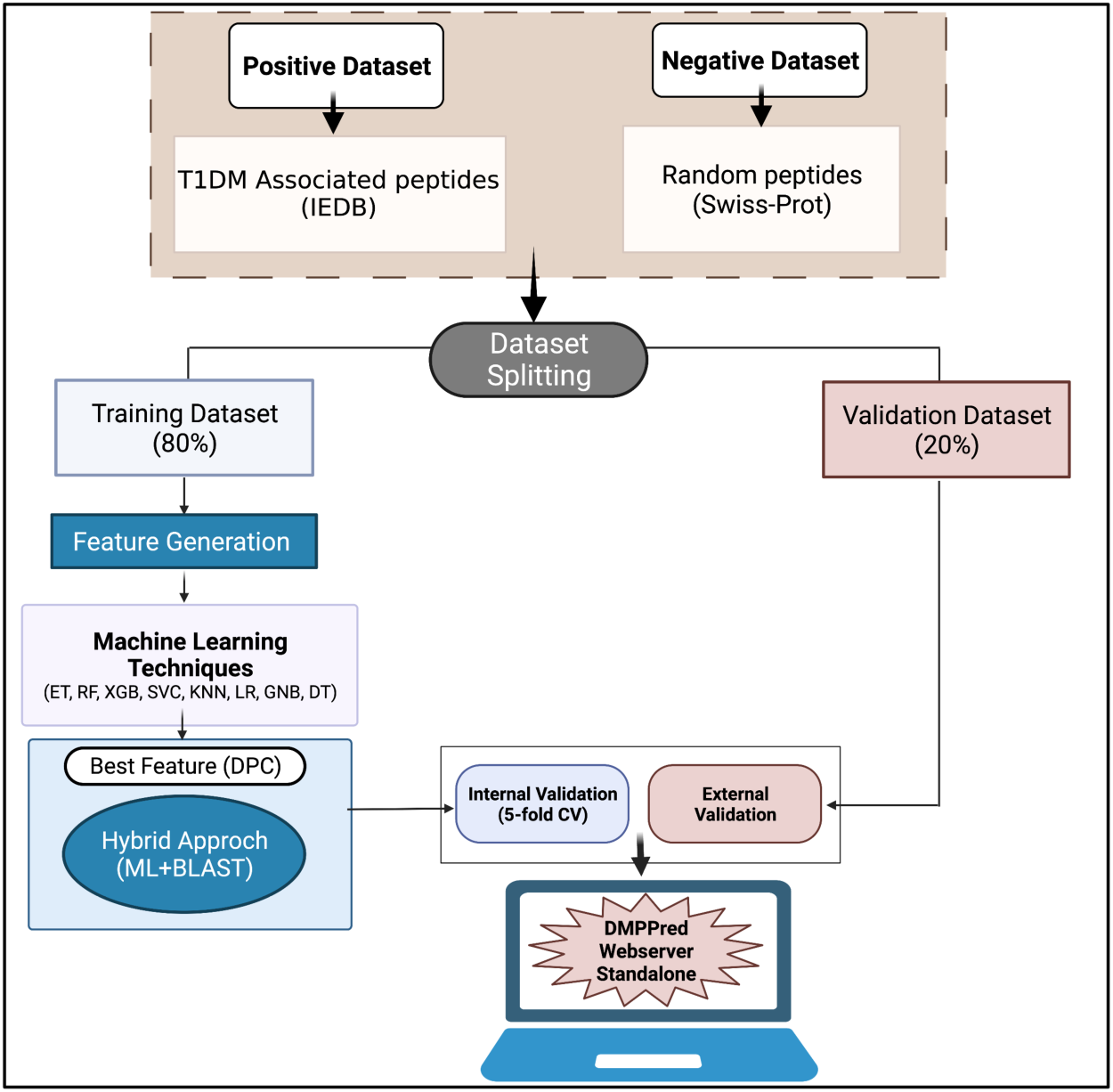
Overall architecture of DMPPred including creation of dataset, training, internal and external validation.

### Positional analysis

In this study, we have developed sequence logo to check the inclination of particular residue at a specific position in T1DM associated peptides. As depicted in Figure 3, the hydrophobic residue Leucine (L) is a highly abundant and conserved residue whereas, Alanine (A) are more preserved at position 2^nd^, 9^th^, 11^th^ and 14^th^ position however, Valine (V) dominates the position 2^nd^, 3^rd^, 9^th^ and 16^th^. On the other side, hydrophilic residue like Glutamic acid (E) is predominant at 2^nd^, 4^th^, 11^th^, and 14^th^ position.

**Figure 3:**
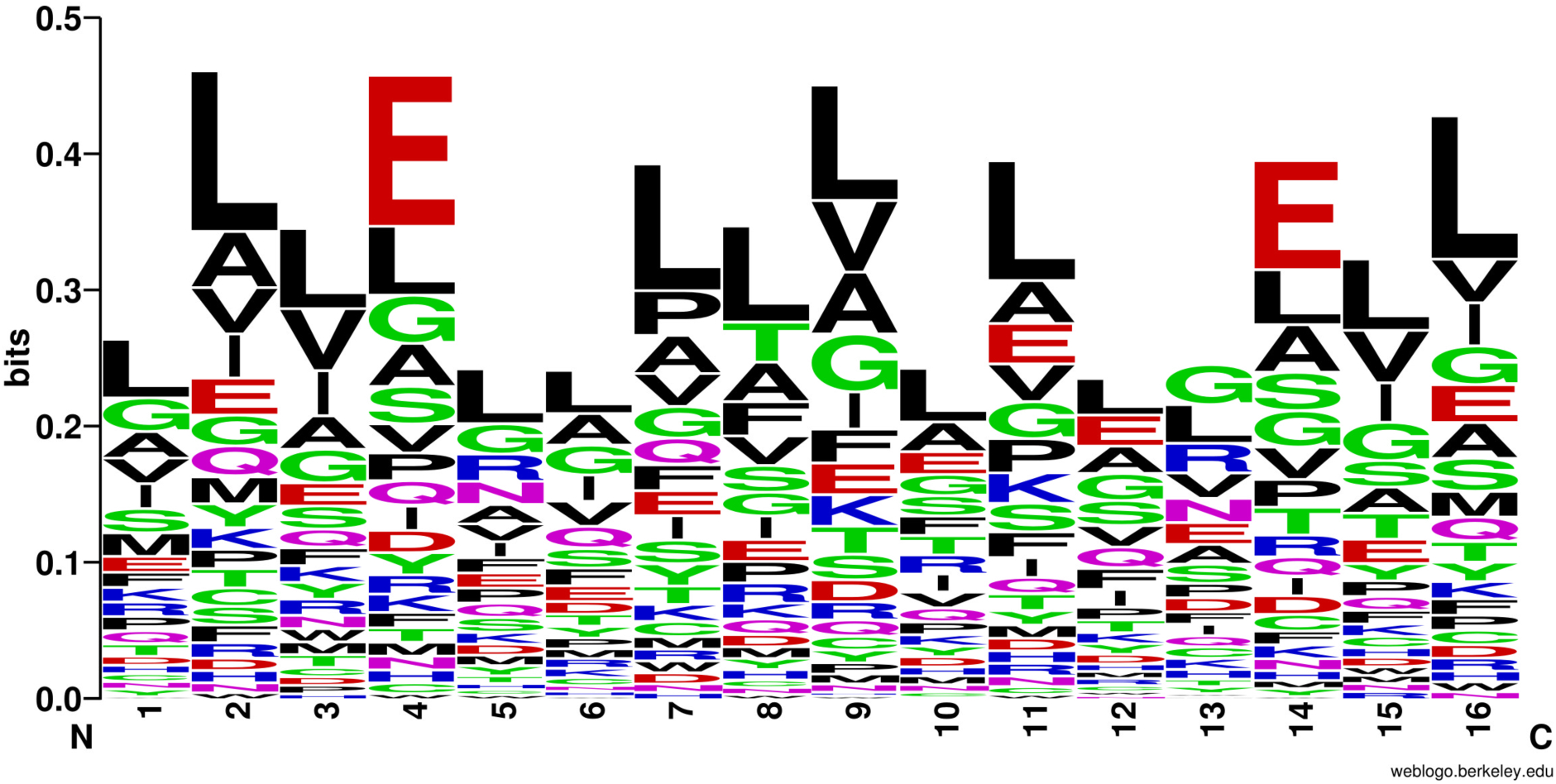
Sequence logo of T1DM associated peptides, Leucine residue is dominated on most of the positions.

### Composition based analysis

Here, we compute the amino-acid composition for T1DM causing and non-causing peptides. Where we calculate average composition of T1DM associated peptides, random peptides and general proteome is shown in Figure 4. The amino-acid residues Leucine (L), Methionine (M), Valine (V) and Glutamic acid (E) are most abundant in the positive dataset, whereas, residues Aspartic acid (D), Lysine (K) and Arginine (R) are highly conserved in the negative dataset.

**Figure 4:**
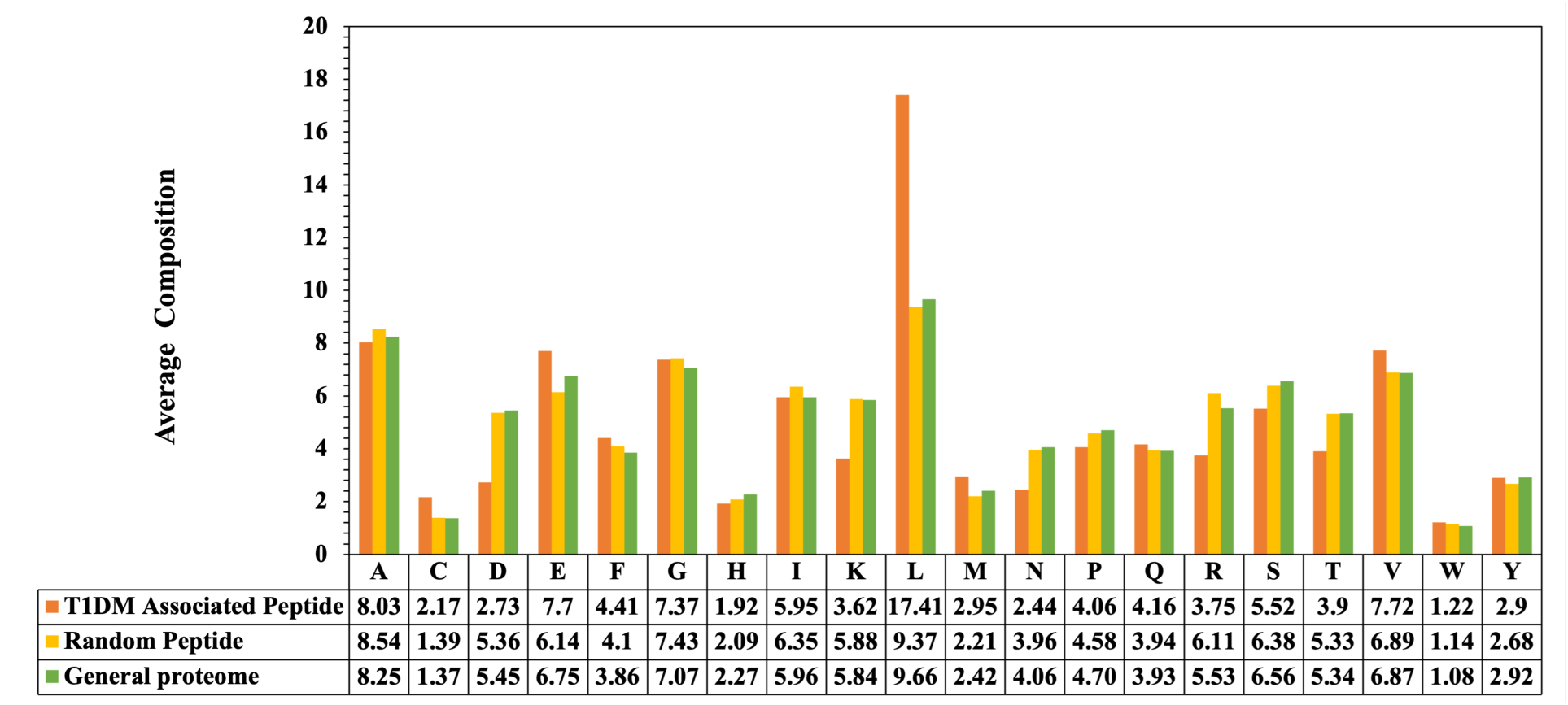
Percent average amino acid composition of T1DM associated peptides, random peptides and general proteome.

### Frequency of HLA-alleles

In the past several studies report that class-II HLA alleles play a key role in the development of T1DM [14]. In order to check the binding efficacy of T1DM associated peptides, we have computed the frequency of HLA-alleles with the T1DM associated peptides. In Table 1, we represented the top 10 class-I and II HLA-binders in the diabetes causing dataset.

**Table 1:** Frequency distribution of HLA-binders in Type 1 diabetes mellitus associated peptides.

| HLA allele | Diabetes causing peptides |
| --- | --- |
| HLA-A*02:01 | 345 |
| HLA-DRB1*04:01 | 185 |
| HLA-DQA1*03:01/DQB1*03:02 | 103 |
| HLA-DR4 | 59 |
| HLA-A2 | 57 |
| HLA-DQA1*05:01/DQB1*03:02 | 45 |
| HLA-DR | 45 |
| HLA-DR3 | 39 |
| HLA-DR53 | 21 |
| HLA-DRB4*01:01 | 20 |

### Machine learning and Performance Evaluation

#### Performance of composition-based features

We used machine-learning algorithms such as DT, RF, LR, GNB, ET, XGB, KNN and SVC in order to develop prediction models. Initially, we developed prediction models based on 12 different types of features computed using Pfeature. We observe that ET-based classifier performs best among all other machine-learning models. In Table 2, we incorporate the prediction performance of all 12 descriptors in terms of the AUROC and MCC. As shown in Table 2, DPC based features outperform other models with an AUROC of 0.893 on training and 0.891 on validation dataset. Models built on AAC based features also perform quite well on both training and validation dataset with an AUROC of 0.869 and 0.870, respectively. Comprehensive results of other classifiers are reported in the Supplementary Table S1.

**Table 2:**
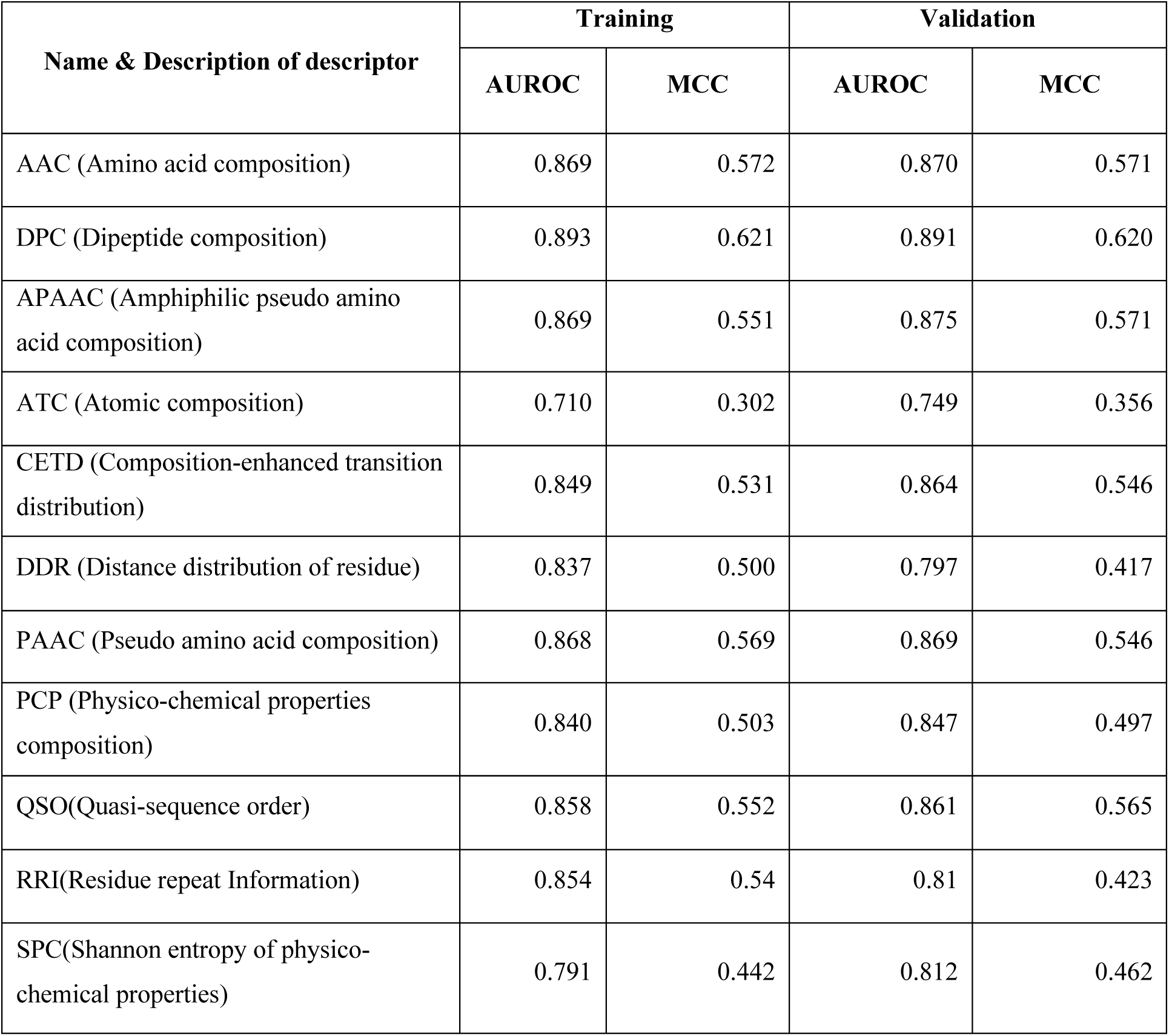
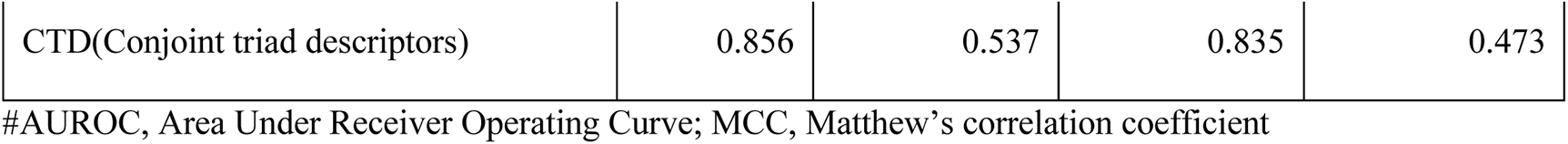
The performance of machine-learning-based models developed on 12 different composition-based features using ET algorithm.

| Name & Description of descriptor | Training |  | Validation |  |
| --- | --- | --- | --- | --- |
|  | AUROC | MCC | AUROC | MCC |
| AAC (Amino acid composition) | 0.869 | 0.572 | 0.870 | 0.571 |
| DPC (Dipeptide composition) | 0.893 | 0.621 | 0.891 | 0.620 |
| APAAC (Amphiphilic pseudo amino acid composition) | 0.869 | 0.551 | 0.875 | 0.571 |
| ATC (Atomic composition) | 0.710 | 0.302 | 0.749 | 0.356 |
| CETD (Composition-enhanced transition distribution) | 0.849 | 0.531 | 0.864 | 0.546 |
| DDR (Distance distribution of residue) | 0.837 | 0.500 | 0.797 | 0.417 |
| PAAC (Pseudo amino acid composition) | 0.868 | 0.569 | 0.869 | 0.546 |
| PCP (Physico-chemical properties composition) | 0.840 | 0.503 | 0.847 | 0.497 |
| QSO(Quasi-sequence order) | 0.858 | 0.552 | 0.861 | 0.565 |
| RRI(Residue repeat Information) | 0.854 | 0.54 | 0.81 | 0.423 |
| SPC(Shannon entropy of physico-chemical properties) | 0.791 | 0.442 | 0.812 | 0.462 |
| CTD(Conjoint triad descriptors) | 0.856 | 0.537 | 0.835 | 0.473 |

#AUROC, Area Under Receiver Operating Curve; MCC, Matthew's correlation coefficient

#### Performance of selected features

As we have obtained maximum performance on DPC features using ET-based models, hence we used these features in order to improve the classification performance. Here, we have used the mRMR feature selection algorithm to obtain the best set of features which can classify T1DM associated peptides and non-T1DM associated peptides. In terms of AUROC, the top-50, 100, and 150 selected features have reasonable discriminatory power. Figure 5, shows the performance (in terms of AUROC) of different classifiers using different feature set for training and validation dataset. The detailed results are provided in the Supplementary Table S2.

**Figure 5:**
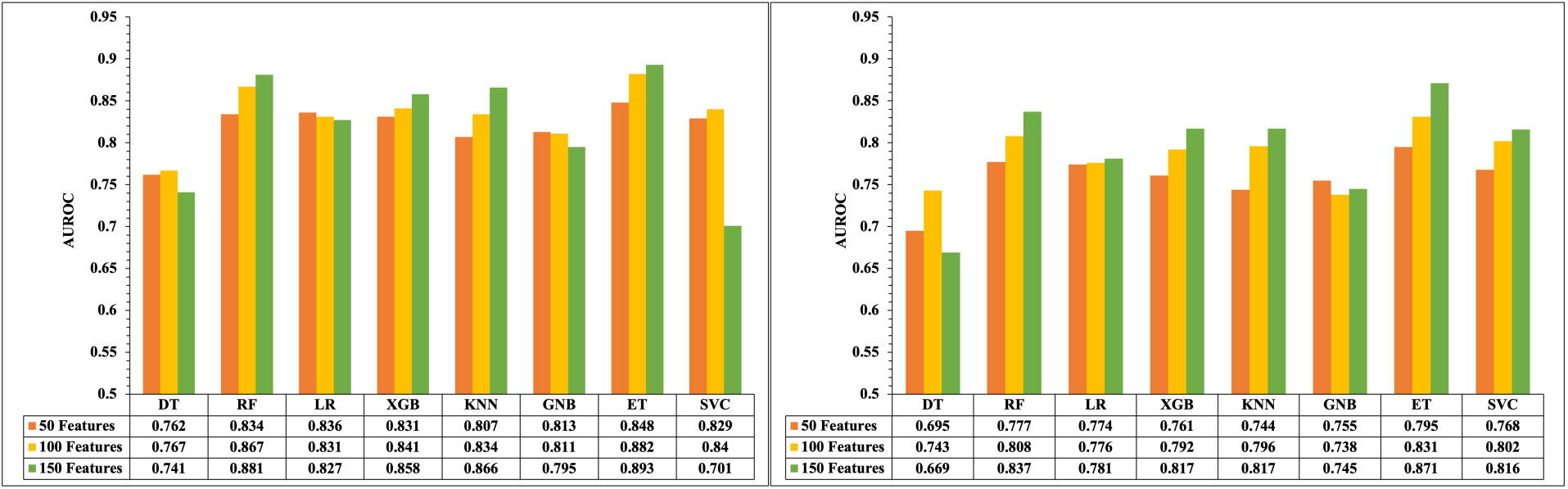
Performance of different classifiers on selected 50, 100 and 150 features.

As shown in Table 3, we have provided the performance of top-150 features selected using mRMR approach. We observed that ET achieves maximum performance with an AUROC 0.893 and 0.871 on training and validation dataset, respectively with balanced sensitivity and specificity. RF-based models also achieve quite similar performance with a slight dip in the accuracy and AUROC on training dataset.

**Table 3:**
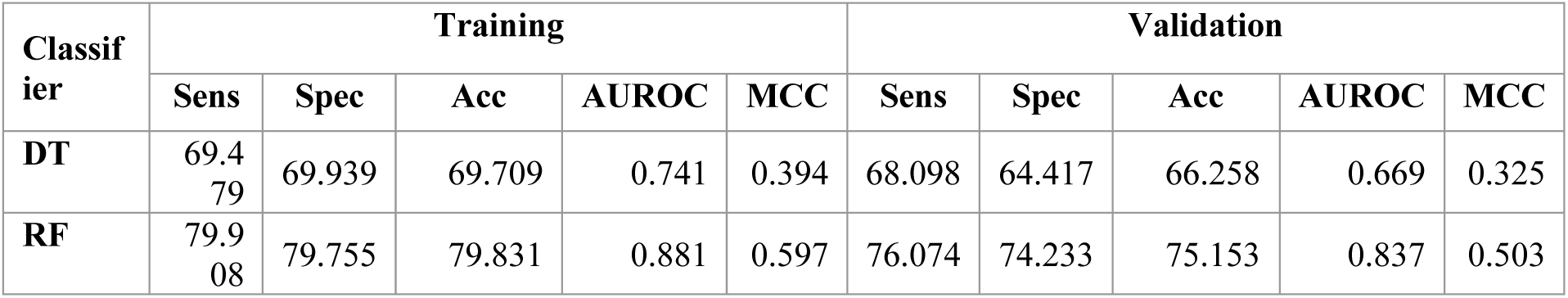
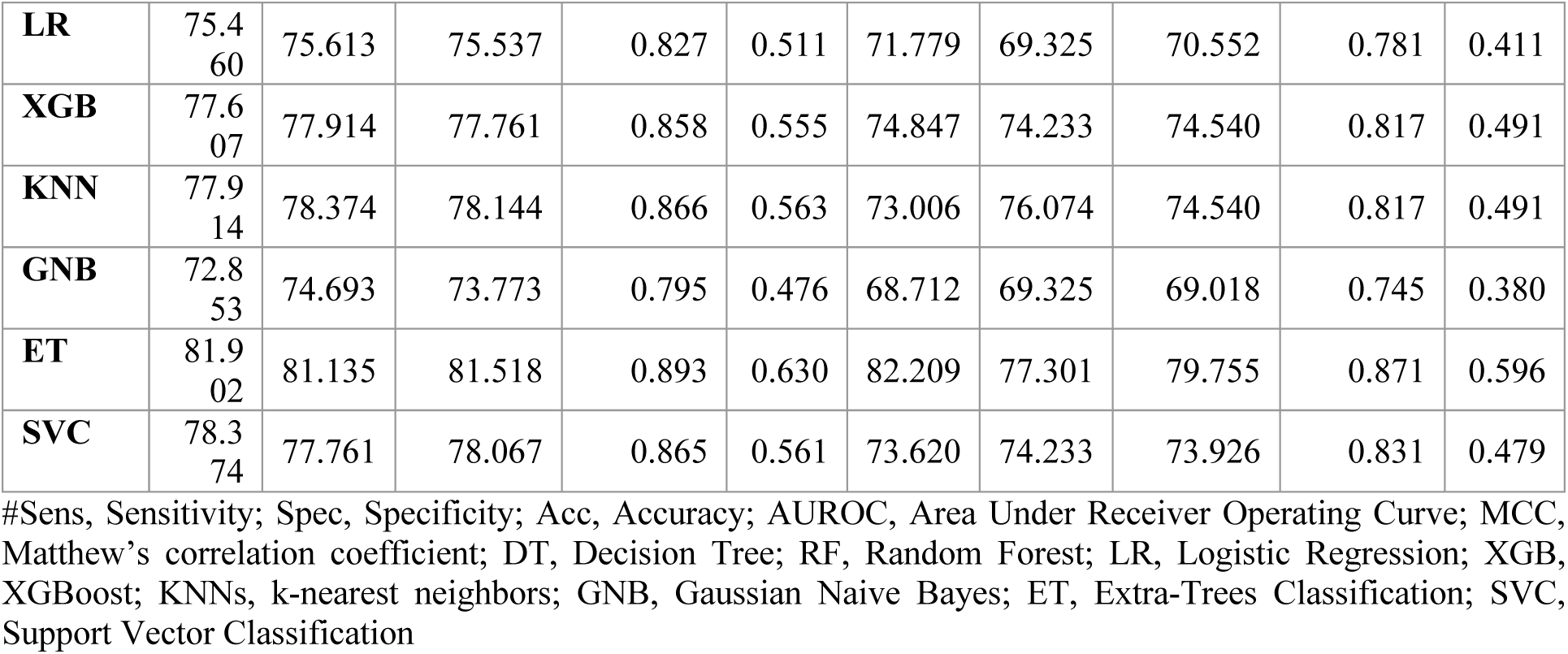
The performance of machine learning-based models developed using DPC-150 selected features on training and validation datasets.

#### Similarity search approach

To further improve our model accuracy, we have designed a similarity search based module based on the BLAST similarity search score, as BLAST is previously used for the annotation and assignment of the functions to a protein on the basis of similarity search [33, 37]. We also used the same approach (blastp) for assigning the given peptide as T1DM associated or non-T1DM associated peptide. We create a local database by using the train dataset against which the query sequence(test set sequences) were searched at an e-value range from 1e^-6^ to 1e^+4^. As shown in Table 4, Probability of correct prediction (chits) ranges from 17.18% - 87.73% on T1DM causing dataset and 0.00% - 61.96% on non-T1DM causing dataset.

**Table 4:** The performance of BLAST-based search on training and validation dataset.

| **E-Value** | **T1DM Causing** |  |  | **Non- T1DM Causing** |  |  |
| --- | --- | --- | --- | --- | --- | --- |
|  | **Chits** | **Whits** | **No-hits** | **Chits** | **Whits** | **No-hits** |
| 1e-6 | 28 (17.18%) | 0 (0.00%) | 135 (82.82%) | 0 (0.00%) | 0 (0.00%) | 163 (100.00%) |
| 1e-5 | 41 (25.15%) | 0 (0.00%) | 122 (74.85%) | 0 (0.00%) | 0 (0.00%) | 163 (100.00%) |
| 1e-4 | 54 (33.13%) | 0 (0.00%) | 109 (66.87%) | 0 (0.00%) | 0 (0.00%) | 163 (100.00%) |
| 1e-3 | 62 (38.04%) | 0 (0.00%) | 101 (61.96%) | 0 (0.00%) | 0 (0.00%) | 163 (100.00%) |
| 1e-2 | 81 (49.69%) | 0 (0.00%) | 82 (50.31%) | 1 (0.61%) | 0 (0.00%) | 162 (99.39%) |
| 1e-1 | 91 (55.83%) | 0 (0.00%) | 72 (44.17%) | 1 (0.61%) | 0 (0.00%) | 162 (99.39%) |
| 1e+0 | 111 (68.10%) | 2 (1.23%) | 50 (30.67%) | 2 (1.23%) | 0 (0.00%) | 161 (98.77%) |
| 1e+1 | 124 (76.07%) | 5 (3.07%) | 34 (20.86%) | 21 (12.88%) | 11 (6.75%) | 131 (80.37%) |
| 1e+2 | 143 (87.73%) | 19 (11.66%) | 1 (0.61%) | 90 (55.21%) | 46 (28.22%) | 27 (16.56%) |
| 2e+2 | 143 (87.73%) | 19 (11.66%) | 1 (0.61%) | 97 (59.51%) | 53 (32.52%) | 13 (7.98%) |
| 1e+3 | 143 (87.73%) | 19 (11.66%) | 1 (0.61%) | 100 (61.35%) | 55 (33.74%) | 8 (4.91%) |
| 1e+4 | 143 (87.73%) | 19 (11.66%) | 1 (0.61%) | 101 (61.96%) | 56 (34.36%) | 6 (3.68%) |

\*Chits: correct hits; Whits: wrong hits

#### Performance of hybrid method

In the current study, we have applied a hybrid/ensemble approach, where we have combined machine learning prediction and BLAST similarity search score. At first a given peptide is classified using BLAST at different e-value. Here we integrated ‘+0.5’ for a correct positive prediction (diabetes causing peptides), ‘-0.5’ for a correct negative prediction (diabetes non-causing peptides) and ‘0’ if hit is not found. This similar approach has been heavily used in previous studies [37, 38]. Secondly the prediction score was computed using ML-based models. Finally, the BLAST score and ML-score integrated, in order to make the prediction as T1DM causing/non-T1DM causing peptide. As shown in Table 2,3 ET-based classifier performs the best using DPC and DPC-150 features. Hence, we generated hybrid models using DPC-150 features (See Table 5). We have calculated performance at different e-values and found that at e-value ‘0.1’ we obtained maximum AUROC of 0.94 and 0.95, with accuracy 88.57 and 90.46 on both training and validation datasets, respectively.

**Table 5:** Model performance developed using Hybrid method (BLAST and DPC-150) on training and validation dataset.

| E-value | Training |  |  |  |  | Validation |  |  |  |  |
| --- | --- | --- | --- | --- | --- | --- | --- | --- | --- | --- |
|  | Sens | Spec | Acc | AUROC | MCC | Sens | Spec | Acc | AUROC | MCC |
| 1e-6 | 83.55 | 83.72 | 83.64 | 0.91 | 0.67 | 81.58 | 90.13 | 85.86 | 0.93 | 0.72 |
| 1e-5 | 84.05 | 84.87 | 84.46 | 0.91 | 0.69 | 81.58 | 91.45 | 86.51 | 0.93 | 0.73 |
| 1e-4 | 84.87 | 84.87 | 84.87 | 0.92 | 0.70 | 82.24 | 91.45 | 86.84 | 0.93 | 0.74 |
| 1e-3 | 85.53 | 85.53 | 85.53 | 0.92 | 0.71 | 84.87 | 91.45 | 88.16 | 0.94 | 0.77 |
| 1e-2 | 86.84 | 86.51 | 86.68 | 0.93 | 0.73 | 85.53 | 92.11 | 88.82 | 0.94 | 0.78 |
| 1e-1 | 88.98 | 88.16 | 88.57 | 0.94 | 0.77 | 87.50 | 93.42 | 90.46 | 0.95 | 0.81 |
| 1e+0 | 89.80 | 89.15 | 89.47 | 0.94 | 0.79 | 86.84 | 90.13 | 88.49 | 0.94 | 0.77 |

#Sens, Sensitivity; Spec, Specificity; Acc, Accuracy; AUROC, Area Under Receiver Operating Characteristics Curve; MCC, Matthews correlation coefficient; ET, Extra-Trees Classification

#### Webserver Interface

To better serve the scientific community, we have developed a user-friendly prediction web interface named ‘DMPPred (https://webs.iiitd.edu.in/raghava/dmppred/) and executed our best models to predict the T1DM associated peptides. The modules available in the web server include “Predict”, “Design”, “Protein Scan” and “Blast Scan”. The module ‘Predict’ allows users to classify the submitted sequence as T1DM associated peptides or random peptides. The ‘Protein Scan’ module allows users to scan or identify T1DM causing regions in the submitted amino-acid sequence. The module ‘Design’ allows users to create all possible analogs of T1DM associated peptides of the input sequence. The module ‘Blast Scan’ allows users to search the query sequence against the database of known T1DM causing peptides. A query sequence is predicted as T1DM causing or random peptide depending upon the match or hit in the database. If found matched or hit in the database predicted as T1DM associated peptide; otherwise random peptide. We also allow users to download the positive and negative dataset that we used in this study available in FASTA file format.

#### Case study

Previous studies shows that there are number of factors responsible for the progression of diabetes. In the past, a number of studies report viruses which promote T1DM including enteroviruses like Coxsackievirus B (CVB) [39], cytomegalovirus [40], mumps virus [41], rotavirus [11, 14, 42-44]. The most commonly found enteroviral strain in diabetic and pre-diabetic patients is CBV4 [45]. We predicted T1DM associated peptides in two proteins of CBV4, polyprotein and VP1 using “Protein Scan” module of DMPpred with peptide length 15 residues. In order to minimize false positive prediction, we used high cut-off score (threshold) 0.70 instead of default cut-off 0.58. Our method predicted 46 T1DM associated peptides in polyprotein which have score 0.70 or more. It was found that none of the peptide have score more than 0.65 in VP1 proteins. It means, polyprotein is dominated by T1DM associated peptides according to our DMPpred. These results agree with existing studies where it has been shown that polyprotein of CBV4 is involved in T1DM [45]. In Table 6, we have shown top 10 highly potential T1DM associated peptides in polyprotein. We have provided the complete results in the Supplementary Table S3 and S4.

**Table 6:** Potential diabetes inducing antigenic peptides predicted by our tool in CBV4 virus polyprotein, selected based on ML score.

| Start | End | Sequence | Hybrid_Score |
| --- | --- | --- | --- |
| 1128 | 1142 | EWLKVKILPEVREKH | 0.97 |
| 1129 | 1143 | WLKVKILPEVREKHE | 0.97 |
| 1130 | 1144 | LKVKILPEVREKHEF | 0.97 |
| 1122 | 1136 | KIQKFIEWLKVKILP | 0.96 |
| 1127 | 1141 | IEWLKVKILPEVREK | 0.92 |
| 1126 | 1140 | FIEWLKVKILPEVRE | 0.89 |
| 1125 | 1139 | KFIEWLKVKILPEVR | 0.86 |
| 1131 | 1145 | KVKILPEVREKHEFL | 0.86 |
| 1123 | 1137 | IQKFIEWLKVKILPE | 0.85 |
| 1124 | 1138 | QKFIEWLKVKILPEV | 0.85 |

## Discussion and Conclusion

The treatment for type 1 diabetes is still a long way off [3]. Currently, only insulin treatment is available that helps in the management of type 1 diabetes [46]. Another therapy used for treatment purposes is glucagon therapy, which is provided at the time when the concentration of blood glucose decreases [47]. A few non-insulin drugs are also available for type 1 diabetes, such as dipeptidyl peptidase-4 inhibitors, glucagon-like peptide-1 receptor agonists, metformin and sodium-glucose co-transporter-2 (SGLT2) inhibitors [48]. Recently developed antigen-specific immunotherapy for T1D have higher efficacy and are safe in clinical outcomes. These antigens are easy to synthesize and can be delivered as proteins, peptides, DNA plasmids or nanoparticles etc. [49-51]. Of note, it is very crucial to identify T1D-specific epitopes/peptides for the experimental and biomedical designing.

In this study, we have proposed a new method for the prediction of T1DM causing and non-T1DM causing peptides. Here we have selected experimentally validated 815 T1DM associated peptides from IEDB. For the negative dataset, we have generated 815 random peptides of the same length using Swiss-Prot database. The compositional and positional analysis reveals that amino-acid ‘L’, ‘M’, and ‘E’ are highly conserved in the T1DM causing peptides in comparison with the negative dataset. We have used Pfeature to compute composition based features using sequence dataset. Here we have computed a total of 1153 features and they were further reduced using mRMR feature selection technique. We have developed various prediction models using different machine learning techniques such as DT, RF, LR, XGB, KNN, GNB, ET and SVC. We observed dipeptide composition based features outperform other models with an AUROC of 0.893 and 0.891 on training and validation dataset. After that, we develop prediction models on selected features i.e., DPC-50, 100 and 150 using mRMR method. DPC-150 features perform best with an AUROC of 0.893 and 0.871 on training and validation dataset using ET classifier. Finally the hybrid model develop by combining BLAST and DPC-150; achieved highest AUROC of 0.945 and 951 on both training and validation dataset. In addition, we have identified the antigenic regions of CVB4 virus, most importantly EWLKVKILPEVREKH, WLKVKILPEVREKHE, LKVKILPEVREKHEF and KIQKFIEWLKVKILP, that induce TIDM with maximum prediction score. We anticipate that, our method has several applications in the field of immunotherapy and vaccine development. As shown in previous studies, antigen based immunotherapy design for specifically targeting T-cell population which can drive disease therefore developing antigen specific T-cell tolerance is required to design for therapeutic use [52, 53]. Our method will provide the facility for the identification of potential disease causing antigens which can cause beta cell destruction in T1DM. In addition, it is also reported that few environmental factors, food, virus and toxins also favours T1DM progression [11, 45, 54-56]. The peptide regions that are associated with the progression of T1DM can be identified using our tool. In addition, several therapeutic peptides have failed during clinical trials due to the presence of allergic and toxin regions. Our method can predict the antigenic regions in therapeutic proteins/peptides that are associated with T1DM. We anticipate that this method serve the scientific community, working in this era. We have provided a user-friendly web-server DMPPred (https://webs.iiitd.edu.in/raghava/dmppred/) for the prediction, scanning and designing of T1DM associated peptides.

## Abbreviations

DM: Diabetes mellitus
T1DM: Type 1 diabetes mellitus
T2DM: Type 2 diabetes mellitus
β-cell: Beta cell
MHC: Major histocompatibility complex
IEDB: Immune Epitope Database
mRMR: Minimum redundancy maximum relevance
HLA: Human leukocyte antigen
CVB: Coxsackievirus B
DT: Decision Tree
RF: Random Forest
SVC: Support Vector Classifier
XGB: XGBoost
LR: Logistic Regression
ET: Extra Tree classifier
KNN: k-Nearest Neighbor
GNB: Gaussian Naive Bayes

## Funding Source

The current work has been supported by the Department of Biotechnology (DBT) funding agency.

## Conflict of interest

The authors declare no competing financial and non-financial interests.

## Authors’ contributions

NK, SP, and GPSR collected and processed the datasets. SP, NK, and GPSR implemented the algorithms and developed the prediction models. AD, SP, NK, and GPSR analysed the results. SP, and SC created the back-end of the web server the front-end user interface. AD, NK, RT, and GPSR penned the manuscript. GPSR conceived and coordinated the project. All authors have read and approved the final manuscript.

## Acknowledgements

Authors are thankful to the University Grants Commission (UGC), Department of Bio-Technology (DBT) and Department of Science and Technology (DST-INSPIRE) for fellowships and financial support, and the Department of Computational Biology, IIITD New Delhi for infrastructure and facilities.

## Data Availability Statement

All the datasets used in this study are available at the “DMPPred” web server, https://webs.iiitd.edu.in/raghava/dmppred/dataset.php.

## Notes

### Competing Interest Statement

The authors have declared no competing interest.

